# Adaptive Nesting Strategies of Dusky Crag Martins in desert ecoregions: Implications for Conservation and Bioinspired Innovations

**DOI:** 10.1101/2024.09.18.613807

**Authors:** Manasi Mukherjee, Aali Pant, Kirti Sankhala, Devavrath Sandeep, Mitali Mukerji

**Author notes:** Corresponding author *Email address:* (Manasi Mukherjee).

## Abstract

Understanding the adaptive strategies of species in extreme environments is crucial for biodiversity conservation. This study is the first to explore the nesting behavior of the Dusky Crag Martin *Ptyonoprogne concolor* in the Thar Desert, Rajasthan. It examined the bird’s site selection, nesting frequency, and construction methods. Soil samples from nest sites and source locations were analyzed to identify structural and elemental properties.

The bird adopts several strategies, such as aligning its nesting period with the monsoon, reusing nests, and utilizing both macro and micro-level additive construction techniques. Detailed analysis revealed a preference for cohesionless sand with low moisture and higher Soil Organic Carbon (SOC) content. The bird strategically alternates light and dark soil layers in its nest. Structural and elemental analyses using optical microscopy and Scanning Electron Microscopy (SEM), and X-ray Fluorescence (XRF), shows the light layers is rich in calcium oxide and fibrous material, and dark layers containing higher iron oxide and partially decomposed litter from *Prosopis cineraria*.

The study concludes that alternating calcium-rich and iron-rich layers enhances energy efficiency, structural integrity, and pathogen resistance in nest construction. his behavior underscores the evolutionary adaptations of *P. concolor* to the extreme desert environment. These findings highlight the importance of conserving desert habitats and provide bio-inspired insights for sustainable building and agricultural practices in arid regions.

## 1. Introduction

Nesting strategies in birds are highly variable and have evolved to enhance survival and reproductive success in diverse habitats and climatic conditions (Perez et al., 2023). These strategies range from simple scrapes on the ground to elaborate structures built in trees or on cliffs. The selection of a nesting site and the construction of the nest are influenced by factors such as predation risk, microclimate, and resource availability, making them crucial adaptations to the environment (Jones, 2001; Kauffman et al., 2021). Birds that build mud nests, such as martins and swallows, exhibit a unique approach to nest construction that is particularly fascinating due to its ecological significance (Piersma, 2013), especially in challenging environments like deserts.

Mud nests are a remarkable adaptation that allows birds to create a stable and protective environment for their offspring in habitats where vegetation may be sparse, and conventional nesting materials are limited (Smith and Smith, 2023). However, the combination of extreme heat, lack of water, and limited cohesive soil presents significant challenges in building materials in desert ecoregions (Amhadi and Assaf, 2019). In such environments, the ability to construct a durable nest with minimal resources is a critical survival strategy.

The Dusky Crag Martin (*Ptyonoprogne concolor*) is an excellent example of a bird that has adapted to these harsh conditions through its nesting behavior. These birds are known for their pure mud nests (Rowley, 1969), which they build on cliffs, rock faces, and man-made structures in arid and semi-arid regions. A striking question arises when considering the nesting behavior of Dusky Crag Martins: How do these birds manage to build sturdy nests from cohesionless sandy soil, especially in deserts where water is scarce and the soil is not naturally cohesive?

Previous studies on mud-nesting martins and swallows have shown that these birds typically select areas near water sources, where they can find cohesive soil (Kilgore and Knudsen, 1977) and use their saliva, rich in mucin (Jung et al., 2021), to bind the materials together. This strategy allows them to construct nests that are both strong and resistant to environmental stressors. However, in water scarce deserts, Dusky Crag Martins may need to employ additional strategies to overcome these challenges.

Our study aims to explore whether Dusky Crag Martins in desert environments have developed unique adaptations to address the extreme conditions they face, such as intense heat, scarcity of water, selective nesting period and the availability of only cohesionless sandy soil. We hypothesize that these birds may have evolved alternative methods for nest construction that are not yet fully understood.

The expected outcomes of this study include a better understanding of the nesting behavior of Dusky Crag Martins, the identification of specific adaptations that allow them to thrive in desert ecosystems, and the potential application of these findings to the development of innovative, bio-inspired building materials and strategies that could be used in challenging environments. These insights will not only enhance our knowledge of avian ecology but also contribute to the growing field of biomimicry, where natural solutions are applied to human challenges.

## 2. Methodology

The study focused on the nesting strategies of *P. concolor* within the desert ecoregion of Indian Institute of technology (IITJ) (26.271386^*°*^ , 73.033335^*°*^), located in the Jodhpur district of Rajasthan. This area lies in Southeastern region of Thar Desert. The campus, predominantly sandy with calcium-rich pockets and abundant native flora, provided an ideal setting for observing the nesting behavior of this species among the four Hirundidae species inhabiting the area.

### 2.1. Nesting behaviour

The nesting behavior and strategies of *P. concolor* were systematically studied over a three-year period, from August 2020 to September 2023. A total of ten nests were monitored to evaluate site selection, nesting frequency, construction methods, and the duration of the nesting process. Each nest was inspected every five days to measure the pace of nest construction and to document the number of individuals involved in the nesting activities. During the study, nesting pairs were tracked to their mud collection sites for site assessment and sample collection. Post-breeding, the abandoned nests were regularly revisited to observe any instances of brood nesting. The study also monitored whether the species returned to the same abandoned nests or the original nesting sites in subsequent breeding seasons. These observations were conducted to gain insights into the adaptive strategies employed by P. concolor in the arid environment.

### 2.2. Characterization and texture analysis

Given that *P. concolor* reuses its nests across seasons, only two nests were carefully removed without disintegration for detailed structure and material analysis, henceforth referred as ‘Nest soil’. This limited removal ensured minimal disruption to the bird population while providing sufficient data for analysis. Based on the observations, soil samples were also collected from the locations where the birds sourced their nesting material, henceforth referred as ‘Source soil’.

#### 2.2.1. Geotechnical analysis

To determine if the soil used by *P. concolor* in nest construction contained silt and clay to enhance cohesiveness, geotechnical analysis was performed on both source and nest soils. The grain size distribution of the soil samples was conducted through sieve analysis. As the soil primarily consists of sand and is non-cohesive in nature, the soil exhibited non-plastic behavior. The liquid limit of the samples was determined using cone penetrometer. The details of the test procedure have been summarized in Table 1.

**Table 1:** Soil Sample and Nest Soil Data.

| Description | Source Soil | Nest Soil |
| --- | --- | --- |
| **Seive Analysis** |  |  |
| Total amount of soil | 300g | 100g |
| **After Washing Soil in 75 Micron Sieve** |  |  |
| Weight of soil retained on 75 micron sieve | 248g | 69.5g |
| Weight of soil passing 75 micron sieve | 50g | 30g |
| **Dry Sieving** |  |  |
| Weight of soil taken for dry sieving | 248g | 69.5g |

#### 2.2.2. Organic Carbon

To investigate if the dark colour of the layer in the nest soil was influenced by organic content, the Soil Organic Carbon (SOC) of both source and nest soils was measured using the Walkley-Black method. This method involved digesting soil samples with 10 ml of 1 N potassium dichromate and 20 ml of concentrated H_2_SO_4_ to oxidize the organic carbon. The reaction mixture was titrated with 0.5 N ferrous ammonium sulfate, using 1 ml of diphenylamine as an indicator, and 10 ml of orthophosphoric acid to enhance the reaction. Four replicates of each sample were analyzed to ensure accuracy, and the average of these replicates was taken as the final reading.

#### 2.2.3. Structural and Elemental characterizations

For the detailed structural analysis and elemental concentration assessment of the source and nest soils, Optical microscopy and Scanning Electron Microscopy (SEM), and X-ray Fluorescence (XRF) methodologies were employed, respectively. Optical microscopy and SEM were utilized to throughly examine the structural characteristics of the soil samples, revealing the structural, compositional and surface morphology. Further, as a particular emphasis was on understanding the key difference between the light and dark layers of a nest, XRF analysis was conducted to determine the elemental composition of the soils. This technique facilitated a precise quantification of the major and trace elements present, such as calcium, iron, magnesium, silica, potassium and titanium. By analyzing the light and dark layers separately, the study could identify variations in elemental concentrations, offering insights into the material selection and adaptive strategies of *P. concolor* in nest construction.

## 3. Results

### 3.1. Nesting behaviour

Findings from 2020 to 2023 showcase the nest-building behavior of *P. con- color* in the Thar Desert. The species initiates nest construction coinciding with the monsoon season, favoring sites with prolonged water retention for soil collection (Fig. 1a). A male-female pair collaborates in this process (Fig. 1b), with each bird making multiple trips (5 to 7 times per visit) to collect soil and regurgitate carefully arranged pellets onto the nest lining (Fig. 1c). It typically takes a two-week period for a pair to create a multilayered nest. All ten observed nests exhibited alternating dark and light soil layers (Fig. 1d). Post construction, the interiors were lined with feathers, dry grass, and *Calotrpis* sp. seed floss (Fig. 1e). Abandoned nests attracted other species like House sparrows and Eurasian collared doves for reuse (Fig. 1f). Remark- ably, the Dusky crag Martins themselves revisited these old nests for future nesting.

**Figure 1:**
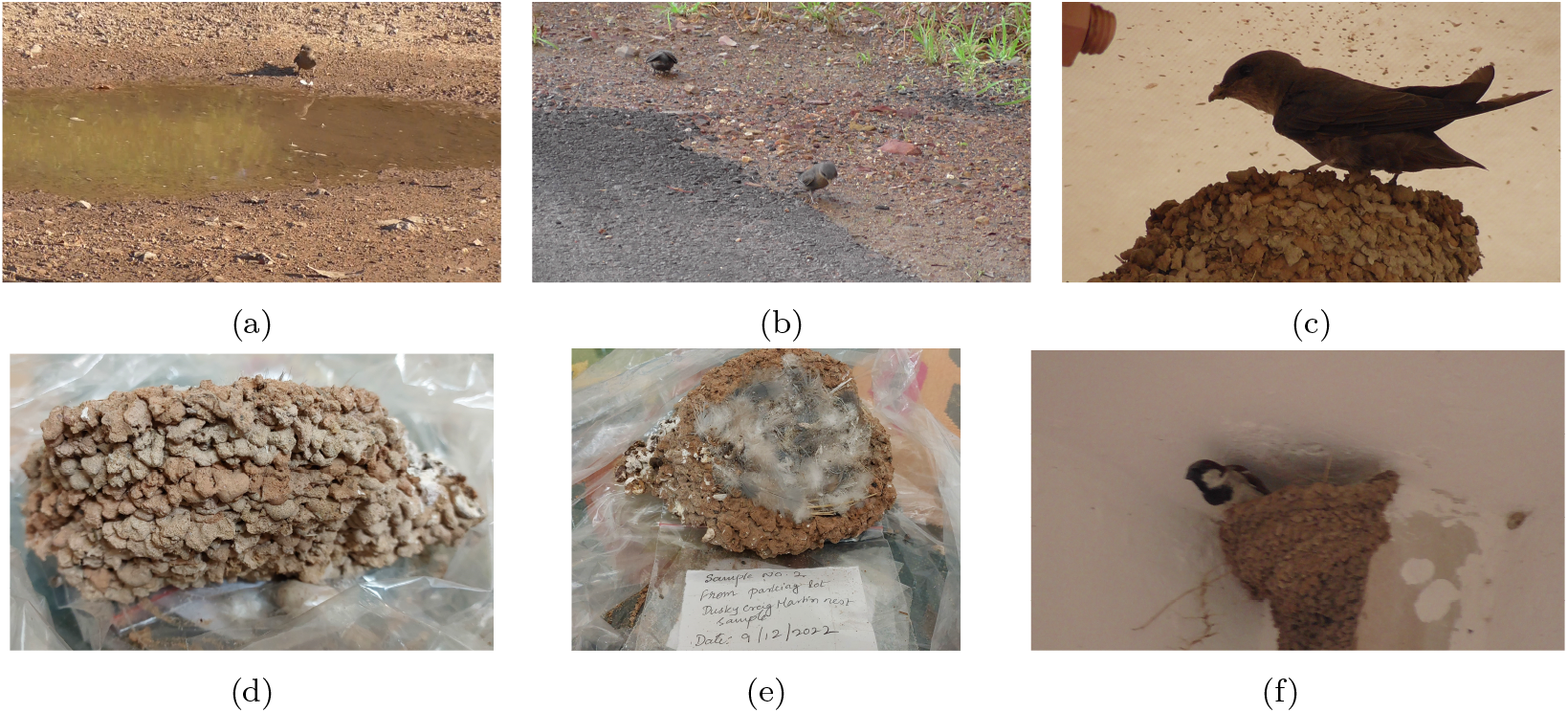
Details of nest making in *P. concolor*. (a) Selection of nesting sites in *P. concolor* based on prolonged water retention. (b) Involvement of a male-female pair in soil collection and (c) nest construction. (d) Notable nest structure displaying alternating light and dark-colored layers. (e) Interior lining of the nest crafted from feathers and *Calotropis* sp. seed floss. (f) Abandoned nests visited by brood birds such as house sparrows.

Nesting sites were meticulously chosen and remained consistent for the species. Seven out of the ten observed nests were established at the entrance vestibules of IITJ campus buildings, while three were situated on parking shade pillars. The half-cup-shaped nests were strategically positioned near the ceiling (Fig. 1f) to minimize predation risks, maintaining maximum height from the floor and minimal available space. Interestingly, on three occasions, the same sites used for prior nests were revisited by the species for subsequent nest construction.

### 3.2. Nest soil characterization and texture

Geo-technical analysis of both source and nest soils revealed that the soil samples consists of predominantly sand sized particles. (Fig. 2). As per the Unified Soil Classification System (USCS) all the soils evaluated in this study can be classified as poorly graded sandy soil (SP). Noteworthy findings indicated no significant differences in basic properties such as gradation curve, specific gravity, and liquid limit between the nest soil and the source soil (Table 2). The primary distinction observed was a 10% increase in finer particles in the nest soils compared to the local soil (Fig.2).Thus the geotechnical analysis results confirmed that both nest and source soil samples were composed of cohesionless sand.

**Table 2:** Geo-technical properties of source and nest soil samples.

| Soil Sample | D50,<br>mm | Cc | Cu | SG | LL<br>(%) |
| --- | --- | --- | --- | --- | --- |
| Soil Sample | 0.14 | 1.05 | 1.88 | 2.75 | 15 |
| Nest Sample1 | 0.2 | 0.79 | 2.94 | 2.59 | 13.7 |
| Nest sample 2 | 0.4 | 0.5 | 7.1 | 2.54 | 12.5 |

**Figure 2:**
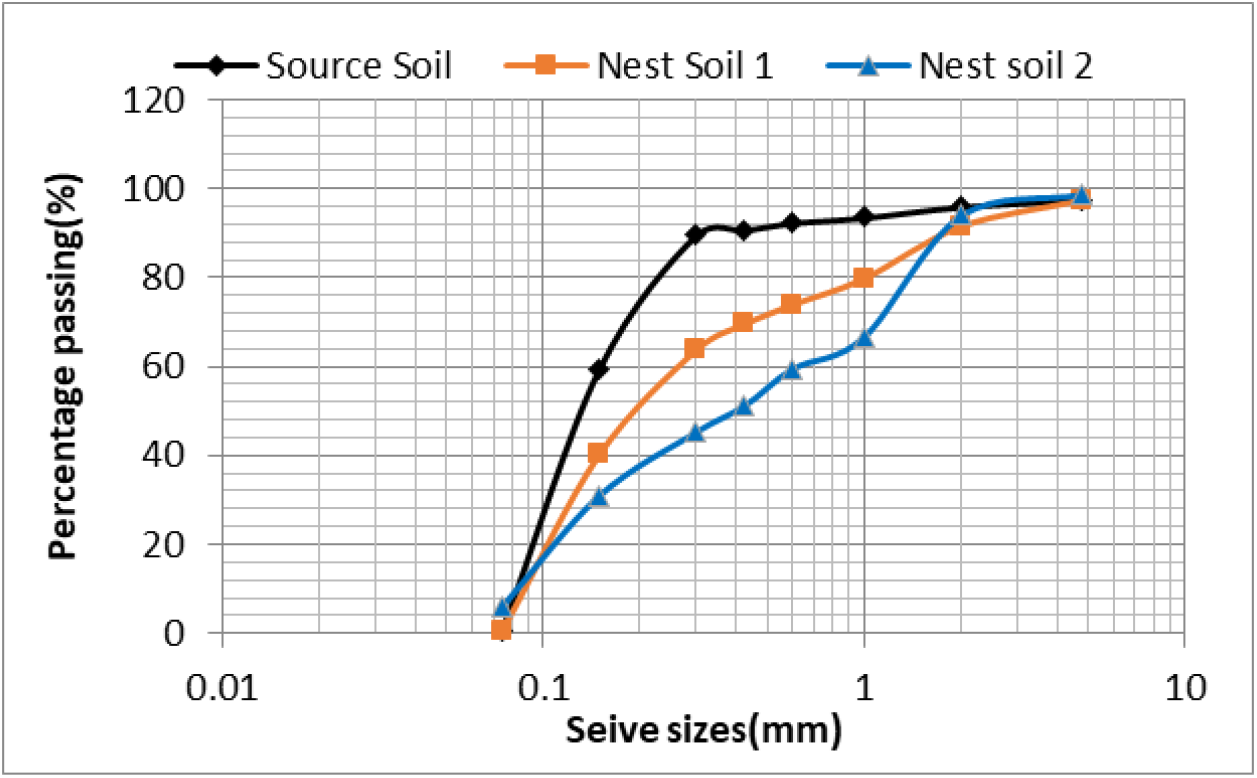
Sieve analysis of source soil and nest soils (Nest Soil 1 and Nest Soil 2) of *P. concolor* to determine the particle size distribution and assess the soil’s mechanical properties. Nest soils exhibit a higher percentage of finer particles compared to the source soil, with approximately a 10% increase in fine particle content.

### 3.3. Nest soil composition and building mechanisms

Furthermore, a closer look at the light and dark layer of the soil reveals the presence of plant matter(Fig. 3a to 3d). To explore whether the binding property exhibited by the nest soil can be attributed to the addition of plant material, the organic carbon content of the soil samples was examined. The mean organic content (%) of the source soil, nest soil 1, and nest soil 2 remained 0.25, 0.35, and 0.48, respectively, indicating a negligible probability of plant matter presence. However, closer examination and optical microscopy revealed difference in the type of plant matter present in the light and dark layers. The light layer showed presence of long, dried, fibrous plant matter (Fig. 3a and 3b), whereas the dark layer was dominated by small, dried leaves and decomposed plant nmatter (Fig. 3c and 3d) .

**Figure 3:**
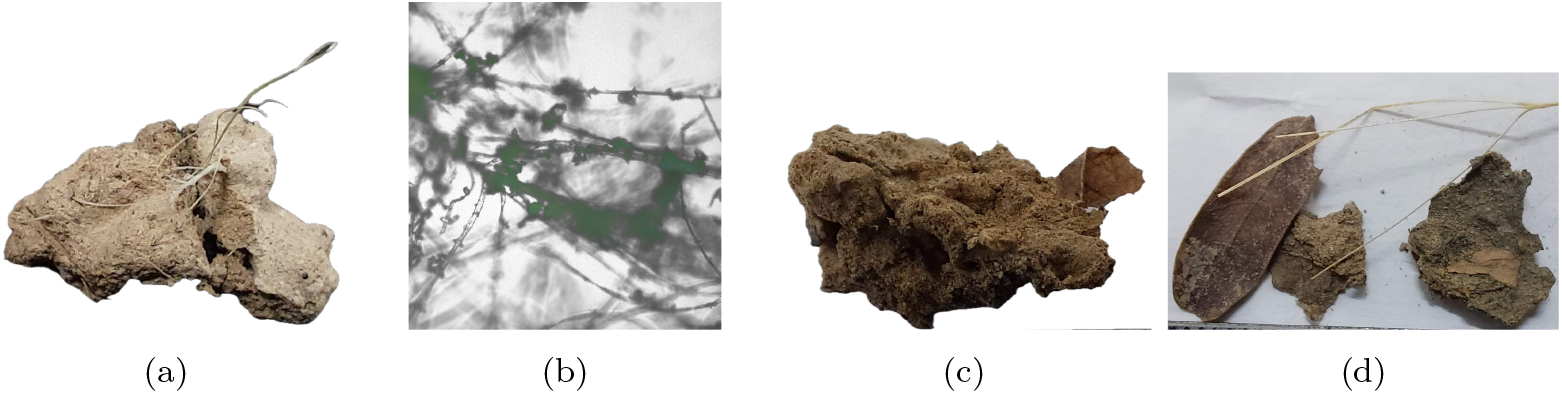
Soil reinforcement strategy used by *P. concolor* ensuring choice of soil enriched with fibrous plant matter for light layer (LL) (a). Fibrous plant matter through bright field optical microscopy (b). The dark layer (DL) consists of large-sized plant litter (c), adjoining the soil particles (d).

As both the physical and chemical properties of the nest soils revealed the absence of cohesiveness needed to build the structure, the possibility of the bird’s additive manufacturing property at micro-scale was analysed. Scanning Electron Microscopy of source soil, light and dark layer of nest samples shows dominance of the sandy nature of the soil (Fig. 4a) that corroborates the results of the geotechnical analysis. Dominance of CaO particles in the light layer is also evident. The elemental concentration of the soils analyzed through X-ray fluorescence (XRF) showed both source and nest soils had dominance of silica (Table 3), constituting more than 50%, which is indicative of the sandy nature of the soil. This was followed by Ca and CaO, which were significantly higher in light when compared to dark layer. Al and Fe and their compounds were higher in the dark layer of the nest soils contributing to the darker colour.

**Table 3:** XRF Analysis of Compounds and Element Concentrations in Source and Nest Soil Layers: Light Layer (LL) and Dark Layer (DL) of DCM.

|  | Compounds concentration (%) |  |  |  |  |  |  | Elements concentration (%) |  |  |  |  |  |  |
| --- | --- | --- | --- | --- | --- | --- | --- | --- | --- | --- | --- | --- | --- | --- |
|  | MgO | Al<sub>2</sub>O<sub>3</sub> | SiO<sub>2</sub> | K<sub>2</sub>O | CaO | TiO<sub>2</sub> | Fe<sub>2</sub>O<sub>3</sub> | Mg | Al | Si | K | Ca | Ti | Fe |
| **SS 1** | 1.8 | 9.7 | 76.1 | 2.9 | 4.4 | 0.5 | 2.5 | 1.6 | 8.3 | 66.6 | 6.2 | 8.5 | 0.8 | 5.3 |
| **SS 2** | 2.4 | 16.0 | 48.8 | 4.4 | 15.2 | 1.0 | 10.7 | 2.1 | 12.6 | 36.5 | 6.8 | 21.7 | 1.3 | 17.0 |
| **NS LL** | 2.4 | 12.4 | 54.9 | 2.8 | 20.7 | 0.8 | 5.4 | 2.1 | 9.6 | 41.3 | 4.6 | 31.0 | 1.1 | 9.2 |
| **NS DL** | 2.5 | 14.8 | 57.7 | 3.3 | 12.7 | 0.9 | 6.7 | 2.2 | 11.9 | 45.4 | 5.6 | 19.9 | 1.2 | 11.8 |

**Figure 4:**
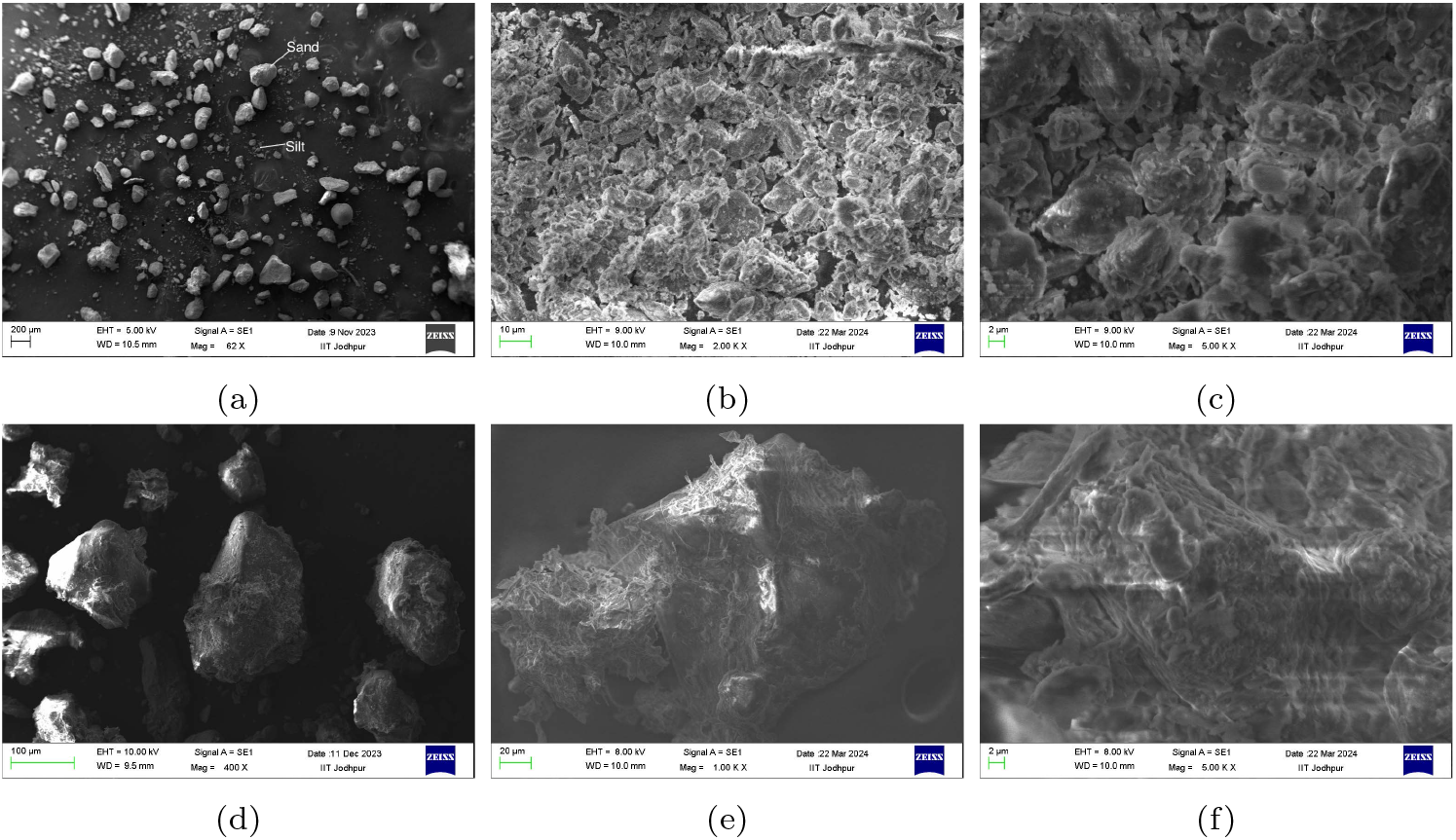
SEM of nest soil samples of *P. concolor*. Panel (a-c) show morphology of sand particles in light layer at different magnifications. (d-f) show morphology of sand particles in light layer at different magnifications

## 4. Discussion

The resources available for *P. concolor* to construct its mud nest in the Thar Desert are limited to poorly graded sandy soil (Fig. 2), minimal vegetative support material, and low moisture content (Table 2). However, the grain size distribution of the soils (Fig. 2) suggest that the bird compensates by selecting soil particles with a broader range of sizes, thereby enhancing the structural strength of the nest (Eyre, 1935). The lower specific gravity of the nest soil samples in comparison to the source soil may be attributed to the presence of organic material (Park et al., 2022), as the nest samples showed higher organic carbon content compared to the source soil. Although the organic carbon content was not indicative of very rich soil (0.48%), it was significantly higher than the mean (0.15%) and approached the maximum value (0.6%) for the Thar desert or arid environments (Moharana et al., 2022), suggesting a relatively organic carbon-rich soil. The primary source of organic carbon in desert soils is the litter fall during the summer, which accumulates in the soil with the onset of rainfall (Liu et al., 2022), coinciding with the nesting season of *P. concolor* in the Thar desert.

Presence of *Prosopis cinereria* leaves and other decomposed material in the dark layer of the nest soil (Fig. 3d) of *P. concolor* indicates the bird’s preference for *P. cinereria* litter-enriched soil. It also indicates the bird’s macro-scale additive manufacturing property in nest construction. Preference to this plant species for food in herbivores (Goyal et al., 1988), for bird nesting (Marathe, 2018) in Thar indicates its larger beneficial properties. *P. cinereria* litter, with its high decomposition rate, organic carbon, nitrogen (N), phosphorus (P), and potassium (K) content, represents the most favorable soil in the arid environment of the Thar desert (Yadav et al., 2008). Furthermore, *P. cinereria* leaves are rich in macrominerals such as calcium and potassium (Sobhy and Al-Rub, 2018), as evidenced by the elemental concentration in the dark layer of the nest soil (Table 3). The anitimicrobial (Bhardwaj, 2021) and antifungal (Joshi et al., 2022) properties of *P. cinereria* leaves offer additional benefits, potentially reducing the risk of disease for eggs and hatchlings.

The alternating use of calcium-rich light layers and iron-rich dark layers in the nesting structure of the Dusky Crag Martin, along with the differences in organic material content, suggests strategic choices by the bird to optimize the nest’s structural and functional properties. While preference for quartz, feldspars and calcite is as established phenomena in many mudnest building birds of Hirundidae family (Papoulis et al., 2018), the use of alternate layers have never been recorded or studied earlier. Additionally, Papoulis et al. (2018) mentions complete absence of any oxides from the nests of all three species of martins and swallows studied. However, our study shows dominance of Calcium and iron oxides from both the source as well as nest soils, varying in percentage composition between layers.

Calcium oxide (CaO) in the light layers (Fig. 4b and 4c) is indicative of the use of calcareous soils with calcium carbonate which is predominant in Thar. Calcium carbonate can enhance the structural integrity of the nest by providing rigidity and strength (Zhang et al., 2021). It has been shown to inhibit the growth of certain pathogens (Ahmed et al., 2023) that might reduce the risk of nest-borne diseases. Larger-sized organic matter is absent in the light layer because it can amplify the reduction in compressive and tensile strength caused by the higher sand content in this layer (Kilgore and Knudsen, 1977). Instead, the fibrous plant material act as a binding material, creating a matrix that holds the soil particles together (Kilgore and Knudsen, 1977), thus preventing disintegration. They also create air pockets, contributing to thermal insulation.

The colour of the dark layer is contributed by higher Fe_2_O_3_ (Table 1) along with the dried leaves (Fig. 3d). The lesser concentration of Calcium in these layers could be balanced by mucin 4e) from the Dusky Crag Martin’s saliva, which could act as a binder. Mucins are known to assemble into a polymer mesh that forms a barrier to molecules and particles while also creating stable, anti-fouling coatings on various surfaces (Ensign et al., 2012; Cone, 2009). Figure 4f indicates mesh formation of the sand particles, binding them together. Mucin thus can bind the sand particles of the nest together, providing additional structural integrity and helping to maintain the nest’s shape and stability. The Fe_2_O_3_ along with mcuin can contribute to moisture retention (Znamenskaya et al., 2013), insulation, protection against harmful microbes (Frenkel and Ribbeck, 2015) and provide a microhabitat that supports beneficial microorganisms (Co et al., 2018).

The birds’ choice of constructing alternating light and dark layers, rather than uniform layers, can be attributed to energy conservation. If salivary mucin were used exclusively as the binding agent, the loose, cohesionless sand prevalent in the source soil would require a larger amount of mucin, thus demanding more energy. Desert birds are known to exhibit physiological adaptations to cope with extreme climates (Dawson and Bartholomew, 1968; Tieleman and Williams, 2000). For these birds, survival involves the challenging tasks of balancing energy and water requirements Williams and Tieleman (2005). Due to the high energy expenditure and water loss associated with the production of mucin, the species has selectively and alternatively used resources like calcium-rich soil and native flora. Timing nest building with the favorable monsoon season also enhances their access to water for both favourable soil (Park et al., 2022) and mucin production for nest building. This nesting behavior thus reflects an evolutionary adaptation to their extreme habitat of deserts, promoting better survival and reproductive outcomes.

Conservation efforts should focus on preserving the natural habitat and plant diversity of the Thar desert to ensure the continued survival of this species. Protecting the desert environment from degradation and invasive species will help maintain the delicate balance required for these birds to thrive.

The nesting strategy of the Dusky Crag Martin offers valuable bio-inspired insights for sustainable solutions in arid environments. Their use of alternating layers with distinct properties for structural integrity, thermal insulation, moisture retention, and pathogen resistance can inspire the design of resilient structures in desert regions (Zhang et al., 2021). For example, incorporating materials that provide both strength and insulation, similar to the bird’s use of calcium and fibrous plant material, can enhance building durability and energy efficiency. Additionally, techniques that improve soil moisture retention and support beneficial microorganisms can boost agricultural productivity in desert soils. Understanding these natural strategies can inform innovative approaches to human challenges in arid landscapes.

## Acknowledgements

The authors would like to thank Dr. Anjali Anand, ICAR-Indian Agricultural Research Institute for helping us with the soil carbon content analysis.

## Authorship contribution statement

**Manasi Mukherjee** contributed in conceptualization, data curation, investigation, methodology, formal analysis, writing original draft, review and editing. **Aali Pant** contributed in methodology, formal analysis, review and editing **Kirti Sankhala** contributed to the methodology, formal analysis, review and editing. **Devavrath Sandeep** contributed in sampling and i **Mitali Mukerji** contributed to supervision, writing-review and editing.

## Data Availability Statement

All data generated or analysed during this study are included in this published article (and its Supplementary Information files).

## Conflict of interest disclosure

The authors declare they have no conflict of interest relating to the content of this article.

## Notes

### Competing Interest Statement

The authors have declared no competing interest.

